# Physical interaction between MSL2 and CLAMP assures direct co-operativity and prevents competition at composite binding sites

**DOI:** 10.1101/2023.04.11.536365

**Authors:** Nikolas Eggers, Fotios Gkountromichos, Silke Krause, Aline Campos-Sparr, Peter B. Becker

## Abstract

MSL2, the DNA-binding subunit of the *Drosophila* dosage compensation complex, cooperates with the ubiquitous protein CLAMP to bind MSL recognition elements (MREs) on the X chromosome. We explore the nature of the cooperative binding to these GA-rich, composite sequence elements in reconstituted naïve embryonic chromatin.

We found that the cooperativity requires physical interaction between both proteins. Remarkably, disruption of this interaction does not lead to indirect, nucleosome-mediated cooperativity as expected, but to competition. The protein interaction apparently not only increases the affinity for composite binding sites, but also locks both proteins in a defined dimeric state that prevents competition.

High Affinity Sites of MSL2 on the X chromosome contain variable numbers of MREs. We find that the cooperation between MSL2/CLAMP is not influenced by MRE clustering or arrangement, but happens largely at the level of individual MREs.

The sites where MSL2/CLAMP bind strongly *in vitro* locate to all chromosomes and show little overlap to an expanded set of X-chromosomal MSL2 *in vivo* binding sites generated by CUT&RUN. Apparently, the intrinsic MSL2/CLAMP cooperativity is limited to a small selection of potential sites *in vivo*. This restriction must be due to components missing in our reconstitution, such as *roX2* lncRNA.

## Introduction

Transcriptional regulation is a complex process involving multiple levels of control. At the first level of the “cis-regulatory code” (1) transcription factors (TFs) bind to specific DNA sequence motifs (2,3). However, in the dynamic and competitive nuclear environment, the affinity of isolated binding sites is insufficient to ensure stable binding. Instead, productive interactions occur at regulatory regions where clusters of binding sites promote TF cooperation and synergistic DNA binding (1). This cooperativity underlies much of the combinatorial regulation of transcription, integrating multiple signaling cues.

To understand the fundamental principles of TF cooperativity and faithful binding site selection, we are studying the *Drosophila* ‘dosage compensation’ system. The male-specific-lethal (MSL) dosage compensation complex (DCC) binds exclusively to the X chromosome in male cells. It boosts the transcription of genes to match the output of the two female X chromosomes (4). The DCC subunit MSL2, the main determinant of X chromosome-selective binding, has only limited intrinsic ability to identify X-specific DNA motifs (5,6). To overcome this limitation, MSL2 cooperates with the ubiquitous zinc finger protein CLAMP (‘chromatin-linked adaptor of MSL proteins’). The two TFs stabilize each other’s binding at some 300 ‘high affinity sites’ (HAS) on the X chromosome (7,8). These sites contain one or more so-called ‘MSL recognition elements’ (MREs), GA-rich motifs that constitute composite TF binding sites (5,9,10).

To study the intrinsic and cooperative DNA binding specificity of MSL2 and CLAMP, we have developed an *in vitro* approach that allows the analysis of TF binding in complex, physiological chromatin at a genome-wide level (11).

The cell-free system relies on extracts from pre-MBT (mid blastula transition) *Drosophila* embryos (DREX, (12)), which assemble maternal stockpiles of chromatin components on genomic DNA. The assembled chromatin shows physiological nucleosome spacing, a rich complement of non-histone proteins and bound insulators surrounded by phased nucleosomes (11,13). In line with its pre-MBT origin, it is transcriptionally inactive and shows dynamic nucleosome mobility catalyzed by abundant ISWI-type nucleosome remodeling complexes. Because endogenous TFs are mostly absent, DREX-reconstituted chromatin provides a naïve substrate to test for genome-wide interactions of purified TFs with thousands of potential binding sites. The experimental strategy mimics the *in vivo* situation at the maternal-to-zygotic transcription transition, when the first TFs encounter a dynamic chromatin (14,15), but additionally allows to manipulate the quantity and quality (mutants) of added TFs at will.

When we explored the ability of MSL2 and CLAMP to bind individual MREs in chromatin we observed strong cooperativity between the two proteins, which was, however, not restricted to functional X chromosomal sites, but included many autosomal sites with similar sequences. We also found that the GAGA factor (GAF) competed with MSL2/CLAMP at non-functional sites, thereby improving the X-chromosomal enrichment (11).

Our current work extends this earlier study in several important ways. First, we investigate the mechanisms that underlie the cooperativity between MSL2 and CLAMP. This cooperativity may involve protein-protein interactions that result in increased residence times of the TFs at composite *cis*-elements and lead to a synergistic effect that exceeds the sum of each individual contribution [direct cooperativity, (2)]. Weak protein-protein interactions may only become relevant when two proteins are brought into close proximity through their binding to adjacent DNA sequence motifs, which is referred to as ‘DNA-mediated’ interaction (16). Interestingly, high-throughput assays that monitor the binding of multiple TFs to a diverse library of sequences have shown that, with the exception of obligatory factor dimerization, the binding of two TFs to neighboring motifs typically leads to additive rather than synergistic effects. This suggests that direct physical cooperation between TFs may not be a common mechanism (17).

In addition to the direct cooperativity of interacting TFs, indirect cooperativity can also be observed in a chromatin context. Nucleosome assembly maximizes the nucleosome organization of genomic DNA (18). TFs often cannot bind to nucleosomal DNA and, therefore, compete with nucleosomes to access their binding sites. At clustered *cis* elements, the successful binding of one TF can hinder nucleosome formation thus facilitating the interaction of other TFs to neighboring DNA. This nucleosome-mediated cooperativity or ‘assisted loading’ of TFs, does not necessarily involve protein interactions and can broaden the combinatorial space (19,20). Nucleosome-mediated cooperativity appears to be the prevalent form of TF cooperativity (21). Some TFs may be better competitors against nucleosomes due to their abundance or mode of DNA interaction, while others may benefit from the action of good competitors (22).

The cooperativity between MSL2 and CLAMP might involve direct protein interactions (8,23). Indeed, we now find that disrupting the physical contact between MSL2 and CLAMP completely abolishes their cooperativity. Surprisingly, rather than the expected nucleosome-mediated cooperativity between non-interacting factors, we find that the stronger GA binder, CLAMP, inhibits the association of MSL2 at shared sites. This suggests that the physical contact between MSL2 and CLAMP not only increases the affinity of MSL2 for target sites (23), but also ensures a defined geometry or registry between the proteins, which not only underlies their cooperation, but also prevents competition. We find that the binding strength of MSL2/CLAMP depends on cooperativity at individual composite binding sites and not on clustering or arrangement of such motifs in the genome.

Finally, we explore to which extent the observed TF cooperativity explains the X chromosome-specific binding of MSL2 *in vivo*. Taking advantage of the superior signal-to-noise ratio and resolution of the CUT&RUN (C&R) technique, we generated the first C&R profiles for MSL2. The new profiles reveal more than twice as many MSL2 binding sites than corresponding Chromatin Immunoprecipitation (ChIP) profiles, which are, remarkably, all on the X chromosome. The improved profiling method not only visualizes HAS, but also degenerate sites of lower affinity that are still restricted to the X (24). Interestingly, we observed only a minimal overlap between the *in vitro* and *in vivo* binding sites. This suggests that the extensive cooperativity between MSL2 and CLAMP observed *in vitro* on all chromosomes is restricted to a relatively small number of functional sites *in vivo*. We discuss a potential role for non-coding *roX* RNA in this process.

## Results

### CLAMP-MSL2 cooperativity is dependent on CLAMP’s N-terminal Zinc finger

We previously mapped interaction surfaces between MSL2 and CLAMP (8). CLAMP binds MSL2 with its N-terminal zinc finger (ZF1, Figure 1A). For MSL2 the interactions required an evolutionary conserved sequence just downstream of the CXC domain, which we termed the CLAMP binding domain (CBD, Figure 1A). Tikhonova *et al*. concluded from NMR studies of isolated domains that the CBD was largely disordered (25). Because unstructured regions are usually not conserved, we used the AlphaFold2 protein structure database to predict the CBD structure. The query of an MSL2 fragment between amino acids 510-690 including both CXC and CBD domains revealed a high-confidence structural prediction for both domains connected by an unstructured linker (Figure 1B). The prediction further showed that CXC and CBD contain surfaces that are compatible with mutual interactions (Figure 1C).

**Figure 1:**
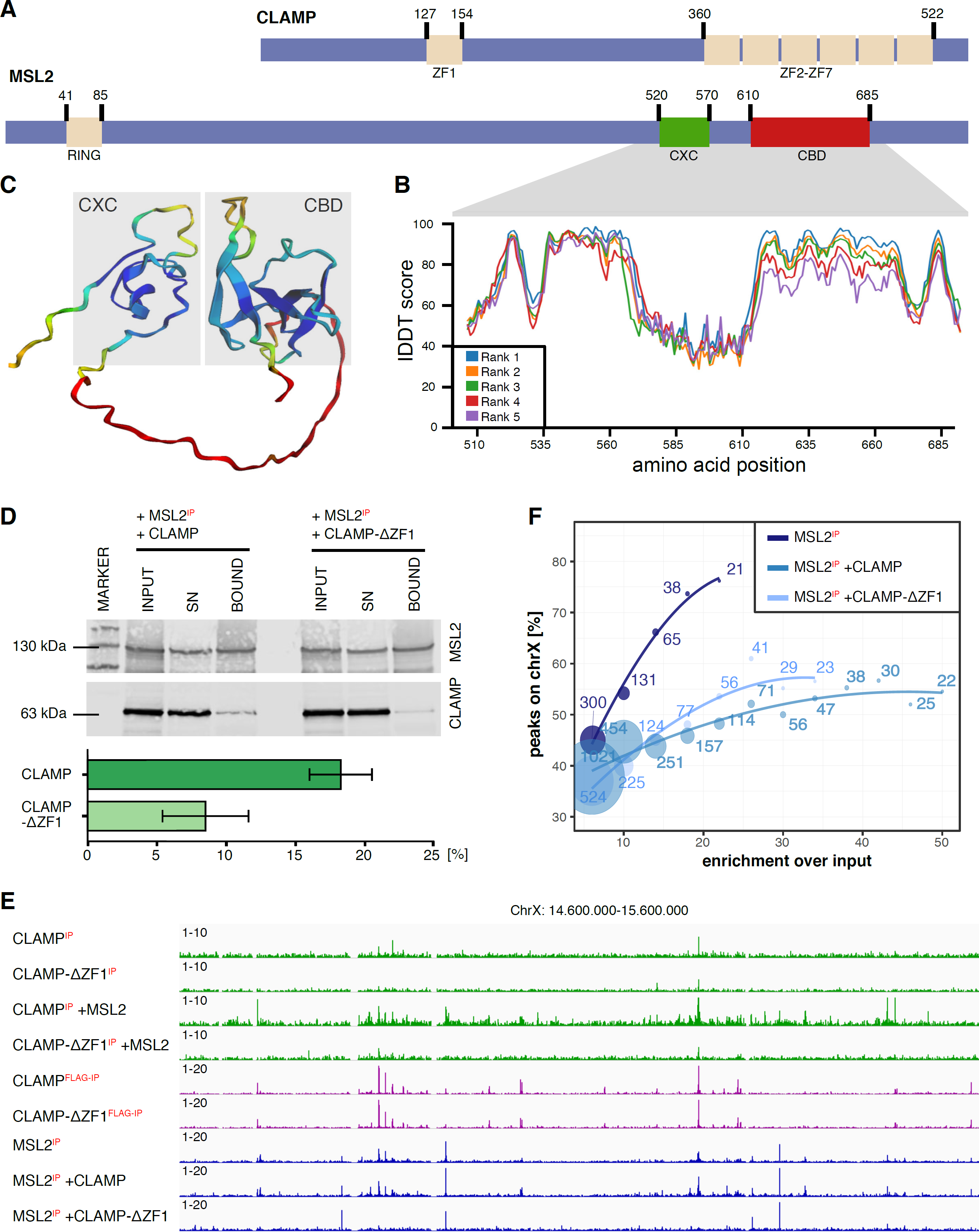
Impact of CLAMP’s first zinc finger on MSL2 binding. (A) The domain organization for CLAMP and MSL2. CLAMP contains seven zinc finger (ZF) domains, of which ZF1 interacts with MSL2 and a subset of ZF2-ZF7 is involved in MRE recognition. MSL2 contains RING, CXC and CLAMP binding domains (CBD). Amino acid positions within the proteins are labeled. (B) Relative IDDT prediction scores from Alphafold2 for MSL2 calculated for amino acids 510-690. High scores indicate high prediction confidence. (C) Predicted protein structure of MSL2 amino acids 510-690, generated by Alphafold2. The IDDT scores from Fig. 1B are color-coded (high=blue, low=red). The CXC and CBD domains are indicated by shading. (D) Western blot analysis of co-immunoprecipitation of MSL2 and CLAMP. 50 nM of MSL2 and either CLAMP or CLAMP-ΔZF1, were added to DREX and immunoprecipitated by anti-MSL2 antibody. The input before IP, supernatant after binding to AB-coupled beads and bound fraction are shown. Blots were probed for MSL2 and CLAMP and imaged using a Licor Odyssey DLx. The amount of coimmunoprecipitated CLAMP was quantified, and the bound fraction was normalized against input amounts and summarized for two sets of protein preparations (see also Figure S1). (E) X chromosome binding profiles. Summarized genome browser profile for all biological replicates shows binding of the protein marked with ‘IP’ determined by ChIP-seq along a representative region of the X chromosome (MSL2-ChIP n=3, CLAMP-ChIP n=2). Maximum values for each window are shown after normalization against the control. For a chromosome-wide view, see Figure S1B. F) X chromosome enrichment of MSL2 peaks in absence and presence of CLAMP or CLAMPΔZF1. Enrichment is related to the respective peak cut-off, defined as the enrichment over background of the peaks. Enrichment is only calculated for datasets with at least 20 remaining peaks.

Attempting to disrupt the interaction between CLAMP and MSL2 we generated protein variants with respective ZF1 and CBD deletions. We found that deleting the CBD yielded a protein that lost all DNA binding activity (not shown), in agreement with earlier findings that partial deletion of the CBD compromises DNA binding (5). In light of the AlphaFold2 prediction it is tempting to speculate that CBD and CXC domains are part of a composite DNA reader module.

Upon deletion of ZF1 (8), CLAMP retained DNA binding activity (see below). When we compared the binding of the wildtype and mutant CLAMP by ChIP, we found the CLAMP-ΔZF1 profile to be systematically weaker (Figure 1E, S1C). We speculated that this may be because the epitope, against which the CLAMP antibody was raised, lies close to the ZF1 deletion site. We therefore generated ChIP-seq profiles for CLAMP and CLAMP-ΔZF1 using an antibody detecting the C-terminal FLAG-tag shared by both proteins. The αFLAG ChIP yielded profiles of roughly similar intensities, indicating that deletion of ZF1 generally does not impair the ability of CLAMP to bind chromatin (Figure 1E, S1C).

To test whether the deletion of ZF1 impairs the interaction with MSL2 in the complex chromatin reconstitution extract, we performed a co-immunoprecipitation assay. MSL2 was incubated with either the wildtype or mutant CLAMP in a chromatin assembly reaction lacking DNA and then immunoprecipitated. The deletion of the ZF1 reduced binding to MSL2 in this crude extract (Figure 1D, S1A).

We then tested the cooperativity of DNA binding of CLAMP and MSL2 in reconstituted chromatin as before (11). Genomic DNA was assembled into chromatin, the TFs were added and crosslinked (Figure S1C). The chromatin was digested with MNase before ChIP and pairedend sequencing of the associated DNA Representative genome browser coverages are shown in Figures 1E and S1B).

In the presence of CLAMP, the binding of MSL2 to chromatin was generally increased, in line with our earlier observations (11). The fact that MSL2 is recruited to strong CLAMP binding sites on autosomes is illustrated in Figure 1F, which relates the strength of binding events (‘signal over input’) to the enrichment on the X. Of the 21 strongest MSL2 peaks in the absence of CLAMP (25-fold over input) more than 75% are located on the X. This illustrates the intrinsic ability of MSL2 to select strong X chromosome determinants. Sites of lower affinity are less enriched on the X (e.g.,131 binding events with 10-fold signal over input reside only 55% on the X). In the presence of CLAMP, MSL2 binds strongly to many more sites, but about half of the strongest sites with 30-50-fold enrichment over input are located on autosomes.

In presence of CLAMP-ΔZF1 the number of MSL2 peaks is reduced and the X-specificity remains low (Figure 1F). This result suggests that ZF1 is required for CLAMP-MSL2 cooperativity and that this cooperativity is direct. Nevertheless, this cooperativity does not explain the exclusive MSL2 binding *in vivo*, a notion that will be further developed, below.

### CLAMP-ΔZF1 competes against MSL2 for MRE binding

To further explore the effect of ZF1 deletion on cooperativity we picked 264 autosomal sites where CLAMP and CLAMP-ΔZF1, when assayed alone, bind strongly as judged by αFLAG IP (Figure 2A, purple tracks). Addition of MSL2 enhanced the binding of CLAMP to these sites, but not of the mutant CLAMP-ΔZF1 (Figure 2A, green tracks). The binding of MSL2 alone was strongly improved by CLAMP. To our surprise, addition of CLAMP-ΔZF1 had the opposite effect: MSL2 binding was reduced (Figure 2A, blue tracks). We had expected that MSL2 would either not be affected by the interaction-defective CLAMP or perhaps even benefit somewhat from indirect, nucleosome-mediated cooperativity. On the contrary, the data suggest that CLAMP-ΔZF1 competes for MSL2 binding. The phenomenon of cooperativity by CLAMP and competition by CLAMP-ΔZF1 was also seen at 100 distinct sites with a PionX sequence motif (Figure 2B)(5).

**Figure 2:**
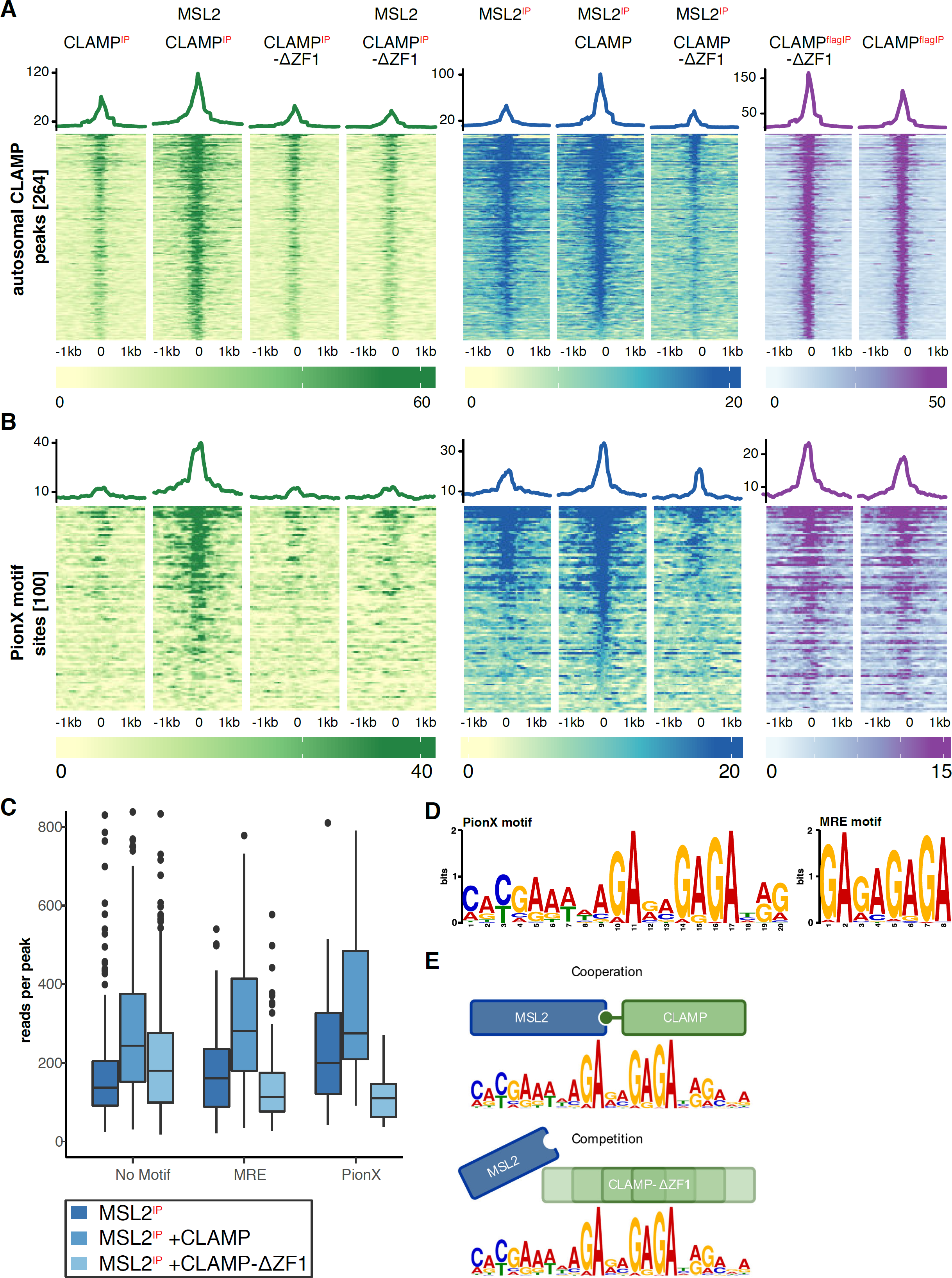
In absence of ZF1, CLAMP competes with MSL2 for binding sites. (A) The annotated proteins were allowed to bind to chromatin-assembled genomic DNA and MSL2 or CLAMP binding determined by ChIP-seq (see Figure S1C). Enrichment of the targeted factor marked with ‘IP’ was plotted to 264 autosomal sites that were bound by both CLAMP and CLAMP-ΔZF1. Heatmaps of individual regions are shown along with average profiles (top). Coverage windows of 2000 bp around the sites were cut out, aligned and the mean for each column calculated. (B) As is A, but plotted at the top 100 PionX motif-containing sites as determined by FIMO search using the PWM in Fig. 2D. (C) Boxplot of normalized read counts per peak. Peaks were determined by Homer for each sample and then separated into groups depending on whether they harbor no strong motif, an MRE motif or a PionX motif, respectively. (D) Position Weight Matrix of the ‘MSL recognition element’ (MRE) and the ‘Pioneering on the X’ (PionX) element used to determine motif-containing sites. (E) Model depicting binding behavior of MSL2 and CLAMP. Intact proteins interact cooperatively and lock into a defined geometry on the large, composite binding motif. In absence of the ZF1 interaction domain CLAMP binds ‘out of phase’ to the motif, leading to competition and displacement of MSL2 from the binding site.

To determine if there was a sequence dependency to the effects seen in the heatmaps we called peaks for MSL2 and divided them into groups according to whether FIMO annotates an MRE, PionX sequence (Figure 2D) or no clear motif. For quantitative assessment we determined the normalised read counts at each peak (Figure 2C). This shows that MSL2 alone binds best at sites with a PionX motif and worst at sites without a clear motif, as expected. Addition of CLAMP increases binding to all sites. CLAMP-ΔZF1 competes with MSL2 only at sites with a clear composite motif. This competition may be due to the fact that MSL2 and CLAMP both recognise GA-dinucleotides in MREs (7,9). At PionX sites, MSL2 may bind the 5’ extension with its CXC domain and more 3’ GA-dinucleotides with parts of the CBD (5). CLAMP-ΔZF1 may compete with this GA-binding. So why does wild-type CLAMP not compete for MSL2 binding? We propose that the direct physical interaction between the two proteins establishes a defined distance between the cooperating proteins, ensuring that they bind to composite sites together and side by side, rather than alone and in competition (Figure 2E).

### Exploring MSL2-CLAMP cooperativity beyond single MREs

The compendium of X-chromosomal binding sites for MSL2 has been redefined several times in recent years. We use a list of 309 HAS defined as overlapping peak regions in two high-quality MSL2 profiles (5). HAS loci often contain multiple MRE and PionX motifs. MSL2 binding affinity may therefore correlate with the number of MREs in a given chromosomal region or with a defined spatial architecture at HAS. This ‘higher order’ level of MSL2 binding as well as MSL2-CLAMP cooperativity has not been systematically investigated so far.

We first explored this question computationally by applying the ‘enhanced chromatin occupancy’ (EChO) strategy (26) to binding data from DREX-assembled genomes. Although this method was developed for CUT&RUN, it can also be applied to ChIP data, which involve MNase digestion and paired-end sequencing. The method exploits the observation that subnucleosome-sized fragments arise from MNase cleavage on either side of a DNA-bound transcription factor. The shortest fragments are centred around a bound TF. When the fragment lengths are plotted across a region of interest, the minima will point to TF contacts on DNA. In absence of binding factors at the sites the profile consists of few reads with lengths of about are 150 bp on average, which reflects the nucleosomal background.

We randomly selected 10 from the 264 autosomal sites where MSL2 and CLAMP bind cooperatively *in vitro* and visualised an area of 401 bp centred on the peaks (Figure 3A). The vertical annotation indicates the positions of the MREs and PionX motifs, revealing a great diversity in the number and arrangement of such sequences. We then plotted the fragment sizes obtained from experiments, in which either CLAMP was assayed alone or MSL2 was mapped in the presence of CLAMP (Figure 3A). The number of reads (individual dots) provides a measure of binding intensity. The minima of the fitted curve points to sites where proteins bind DNA. Several conclusions can be drawn from the plots. (1) The predicted TF binding sites generally match the annotated MREs and PionX sequences. (2) Strong binding events often coincide with MRE clustering, but there are also cases where binding occurs in the absence of annotated MRE motifs. (3) The curves for CLAMP alone and MSL2 in the presence of CLAMP generally, but not always, follow a similar path.

**Figure 3:**
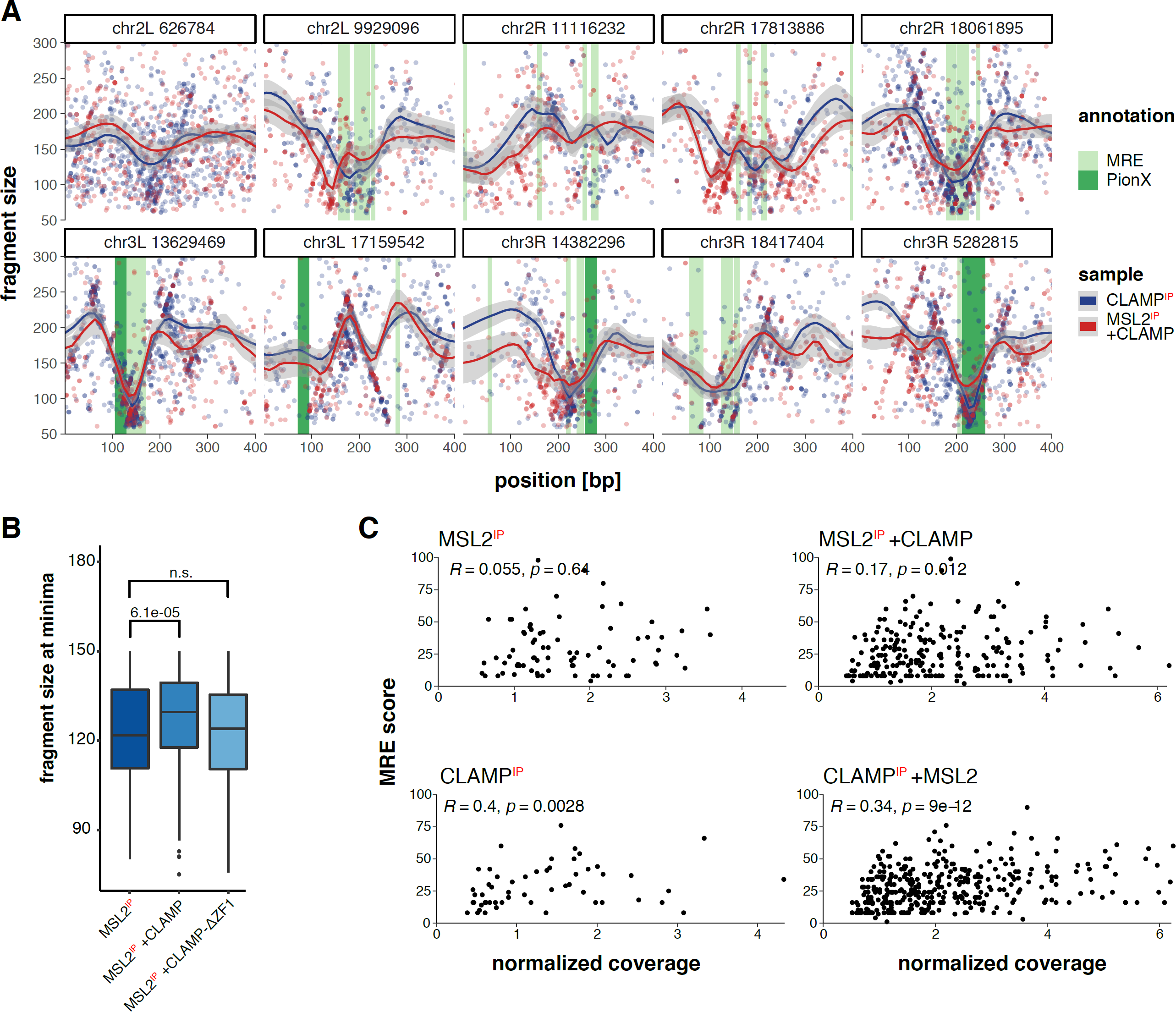
MSL2 binding sites show heterogeneous motif occurrence and binding behavior. (A) MSL2 and CLAMP binding was determined by ChIP-seq after chromatin assembly. Ten binding sites bound by both proteins were sampled from the sites shown in Fig. 2A. The size of individual reads within the binding sites was plotted against their relative position. MRE and PionX sites were determined using the PWMs from Fig. 2D and are annotated with green shading. A regression curve is fitted using loess and the shaded area represents the 5-95% confidence interval. As TFs protect a smaller area of DNA from cleavage then nucleosomes, the precise binding site of a TF can be found by determining the minima of fragment sizes within a given peak. (B) Using the regression curves the minima within each peak are determined and the size of all minima <150 bp are summarized. Boxplots are drawn and the correlation between groups measured using Wilcoxon signed-rank test. P values are annotated above the boxes. (C) Reads per peak were normalized to total read count and plotted against the MRE score. MRE score was determined as the number of GA dinucleotides that fall within MREs, as defined by a FIMO search using the PWM shown in Fig. 2D.

The data suggest that CLAMP and MSL2 do in fact bind together to the composite sequence elements. If this was the case, an increased MNase ‘footprint’ (indicated by increased fragment sizes) would be detectable at sites where more than one TF binds. We computed the fragments lengths across all peak areas in all *in vitro* binding experiments, determined minima at each site by loess regression and summarized all the reads found at these locations (Figure 3B). The ‘fragment size at minima’ gives a relative measure of the TF footprint. MSL2 binding alone correlated with an average fragment length of 120 bp. In the presence of CLAMP, the average length of fragments was increased by 10 bp. In contrast, addition of CLAMP-ΔZF1 had no effect. These findings support the notion of cooperative binding of MSL2 and CLAMP at individual composite binding sites, which are largely defined by the presence of CLAMP.

The anecdotal fragment size profiles (Figure 3A) illustrate a great diversity of binding site architecture, but they do not allow general conclusions about binding cooperativity between clustered MRE sequences. To address this question more systematically, we determined an ‘MRE score’ for each peak area (see methods) by summing up all nucleotides contained in an 8 bp MRE PWM (note that PionX sequences conform with the general MRE definition). The higher the MRE score, the more MRE motifs cluster at a given site. We then related this MRE score to the corresponding binding strength, indicated by the normalized read coverage (Figure 3C). For the binding of MSL2 alone, the correlation between MRE score and binding strength was poor. This is explained by the fact that MSL2 alone binds best to single PionX sequences and not so much to extended GA-rich regions (5,11). In contrast, CLAMP alone is known to bind extended GA sequences (27), which is reflected by a good correlation between MRE score and binding strength. If MSL2 binding strength is assessed in the presence of CLAMP, the correlation to MRE score remains poor (although slightly better than for MSL2 alone). Interestingly, there are many sites with a single MRE (MRE score of 8) with very different binding strengths. Finally, if CLAMP binding is assessed in the presence of MSL2 the correlation to MRE score is reduced, presumably because MSL2 can stabilize CLAMP at otherwise low affinity binding sites (Figure 2B). We conclude that the cooperativity in chromatin binding between MSL2 and CLAMP occurs predominantly at the level of individual, composite binding sites and less so between TFs bound to clustered elements within a HAS.

### CUT&RUN detects X chromosomal MSL2 binding sites beyond the known HAS

These findings, along with our previous *in vitro* genome-wide binding assays (8,11), reveal fundamental principles of cooperativity and competition at GA-rich MRE sequences on all chromosomes. However, *in vivo* the cooperativity between MSL2 and CLAMP is strictly limited to X chromosomal HAS (8). This discrepancy may be resolved *in vivo*, if the cooperativity at the level of MRE clustering was more important, due to chromosome organization (28).

To obtain binding profiles suitable for EChO analyses, we generated the first C&R profiles for MSL2 in the male S2 cell line. The C&R methodology avoids immunoprecipitation and thus yields a better signal-to-noise ratio and improved resolution (29). It also avoids formaldehyde crosslinking, a major source of inconsistency between protocols. If combined with paired-end sequencing, C&R is ideal for EChO plot analysis.

Our C&R analysis impressively documented the exclusive binding of MSL2 to the X chromosome *in vivo*, and in this respect was superior to previous ChIP-seq profiles [Figure 4A; (5)]. C&R yielded twice as many MSL2 binding sites (709) than corresponding ChIP profiles. Apparently, the improved profiling method not only visualizes HAS, but also degenerate sites of lower affinity that are nevertheless restricted to the X (24,30). As such sites have not been previously characterized, we divided the total set of peaks into thirds according to their binding strength and performed motif searches using MEME. The ‘top’ peaks, where MSL2 binds most strongly, contain sequences that resemble the ideal PionX motif (Figure 4B). Binding sites of intermediate affinity yield the familiar MRE motif in a longer GA-rich context. The ‘bottom’ group of weaker MSL2 interaction contains more degenerate motifs that are still GA-rich. Even these low-affinity motifs tend to avoid the minimal binding site of GAF, GAGAG. We previously showed that GAF can outcompete MSL2/CLAMP at sites containing GAGAG sequences, thereby excluding non-functional sites from MSL2 interaction (11). TF binding motifs are typically short, often less than 10 bp. The fact that MEME detects PWM signatures that are 20 bp and longer (Figure 4B) supports the notion that cooperation and competition between MSL2 and CLAMP occur at single, long composite binding sites.

**Figure 4:**
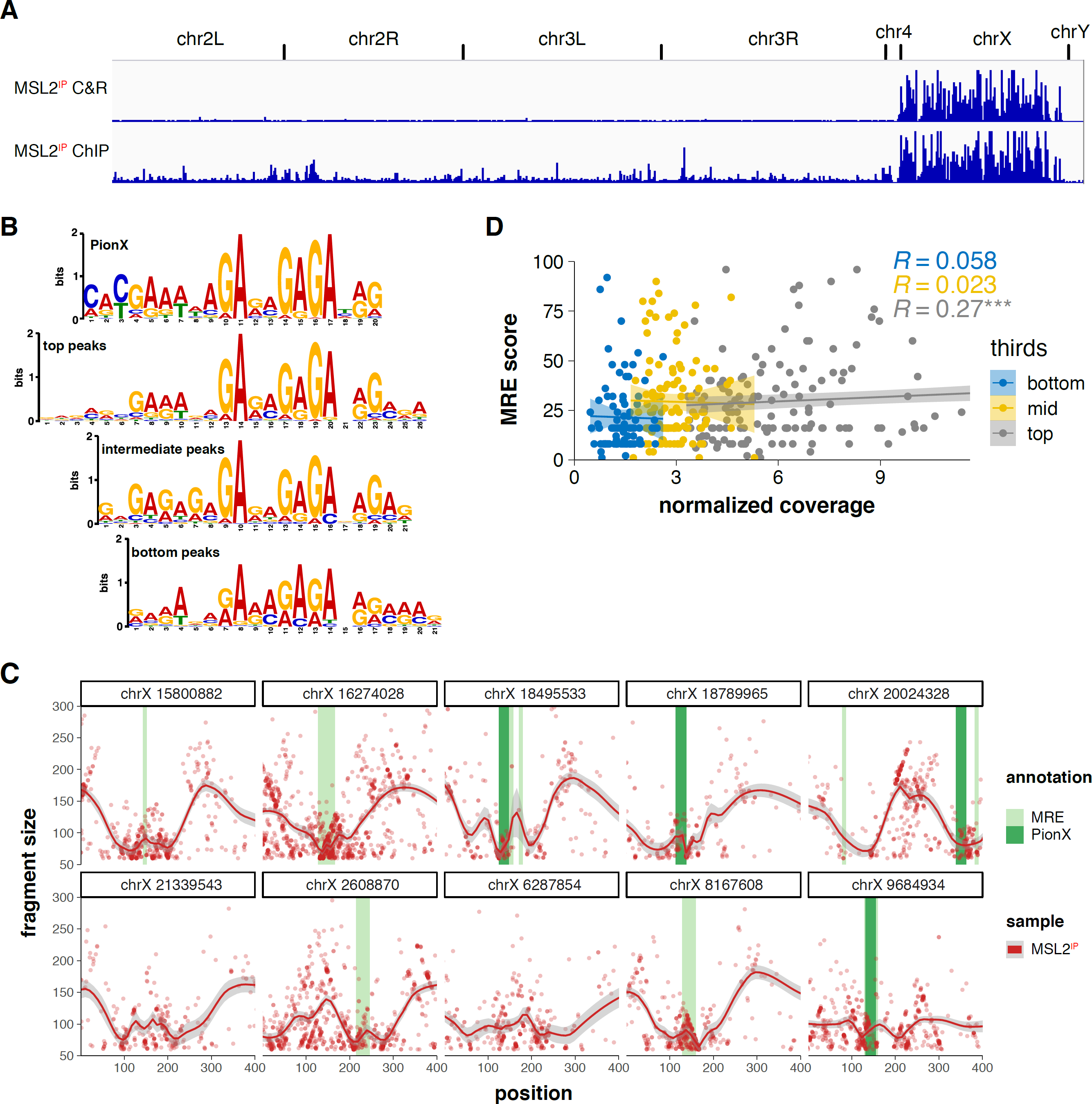
An MSL2 CUT&RUN (C&R) analysis allows insights into motif and binding architecture within peaks *in vivo*. (A) Genome browser profile over the *Drosophila* genome summed up from three biological replicates of C&R profiles for MSL2 in S2 cells (‘C&R’) or ChIP-seq (‘ChIP’) ((5), n=3). Maximum values for each window are shown and chromosomes are annotated. (B) Position Weight Matrix (PWM) of the PionX motif compared to the PWMs found by MSL2 C&R peaks as determined by MEME. The 709 peaks were split into thirds depending on their enrichment over input and analyzed separately. (C) MSL2 binding was determined by C&R *in vivo*. Peaks were determined by Homer. For each peak the fragment length of individual reads was plotted against their respective position within the peak. Ten binding sites were sampled from the top third of binding sites and are used as illustration. MRE and PionX sites were determined using the PWMs from Fig. 2D and are annotated. A regression curve is fitted using loess and the shaded area represents the 595% confidence interval. As TFs protect a smaller area of DNA from cleavage then nucleosomes, the precise binding site of a TF can be found by determining the minima of fragment sizes within a given peak. (D) Reads per peak were normalized to total read count and plotted against the MRE score. MRE score was determined as the amount of GAs that fall within an MRE. MREs were found using a FIMO search using the PWM shown in Fig. 2D. The 709 peaks found by MSL2 C&R were split into thirds depending on their enrichment over input and analyzed separately.

Looking at the clustering of MRE motifs at MSL2 binding sites, we again observe a large diversity in the number and arrangement of MRE, similar to the *in vitro* sites. Figure 4C shows 10 randomly sampled sites for which the motif positions were determined by FIMO, as before. For EChO analysis, we plotted the lengths of individual reads within each binding site. The minima of the fitted curve indicate protein binding sites. Unlike the *in vitro* experiment, where we can control the input of TFs, we do not know which other proteins bind along with MSL2 at these sites. The fact that we sometimes observe more than one minimum and that some minima are broad suggests that other factors, such as CLAMP, bind in the vicinity of MSL2. To approach the question of inter-MRE cooperativity more quantitatively, we calculated the MRE score within each site and related it to the binding strength, as before. For this evaluation of *in vivo* sites, the ‘top’, intermediate and ‘bottom’ thirds of the peak areas were analysed separately (Figure 4D). Linear regression shows no correlation for the two lower thirds of MSL2 peaks *in vivo* (0.058 and 0.023 respectively) and a very weak correlation for the upper third of MSL2 peaks (0.27). This finding suggests that, similar to the *in vitro* analysis, the binding strength is mainly determined by the interaction of MSL2 with individual MRE motifs and not dependent on the number or arrangement of MRE-type sequences.

### The intrinsic DNA binding cooperativity between MSL2 and CLAMP is largely overridden in vivo

So far, the *in vitro* and *in vivo* analyses have yielded MSL2 binding sites that appear similarly diverse with respect to number, MRE PWM and arrangement. We therefore expected the *in vitro* sites to overlap considerably with the expanded set of X chromosome sites obtained by C&R. To our surprise, the two sets of binding sites were very distinct. Heat maps sorted according to the strength of the 709 binding sites identified by C&R reveals that only a small fraction of these sites is actually bound *in vitro* (Figure 5A). This is best illustrated by the Venn diagram in Figure 5B. The MSL2 C&R peaks overlap only minimally with the sites bound by MSL2 in the presence of CLAMP. We had previously shown that GAF plays an important role in occluding non-functional MSL2-CLAMP binding sites *in vitro* (11). Indeed, about one third (35/111) of the *in vitro* peaks in presence of CLAMP and GAF overlap with *in vivo* C&R sites. We conclude that the *in vivo* binding profile cannot be explained by intrinsic DNA binding cooperativity of MSL2 and CLAMP, but that this cooperativity is largely offset *in vivo* and only allowed in a context-specific manner.

**Figure 5:**
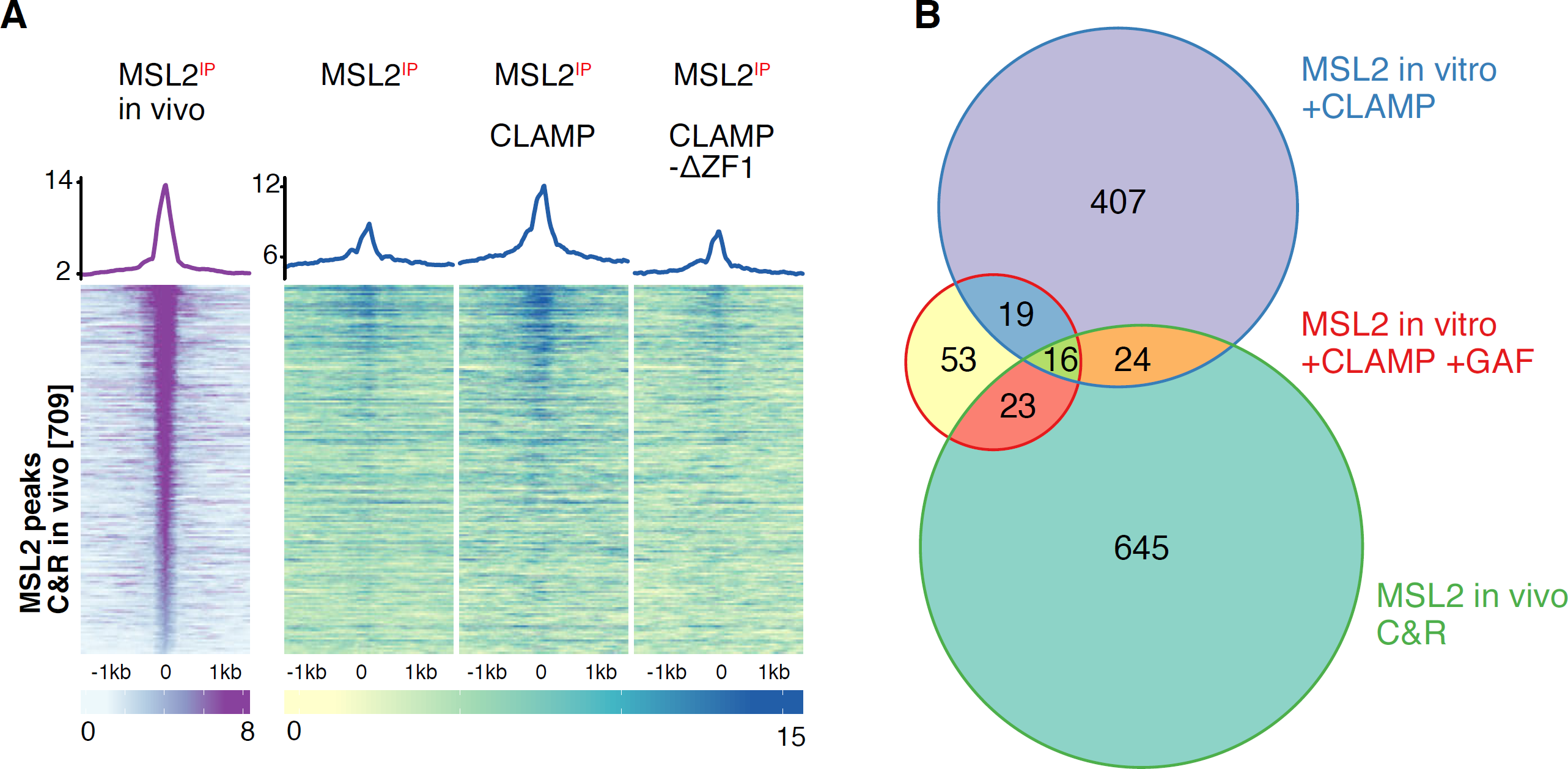
The intrinsic MSL2-CLAMP cooperativity found *in vitro*, is restricted *in vivo* by other factors. (A) MSL2 binding was determined by C&R *in vivo* and by ChIP-seq *in vitro*. Enrichment was plotted to 709 MSL2 C&R sites and illustrated by average profiles (top) and heatmaps of individual regions. Coverage windows of 2000 bp around the sites were cut out, aligned and the mean for each column calculated. (B) Venn Diagram of MSL2 peaks found by C&R *in vivo* and by ChIP-seq *in vitro*. MSL2 *in vitro* peaks in presence of CLAMP and GAF were taken from (11).

## Discussion

Our experimental strategy involves the assembly of physiological chromatin on genomic DNA to provide a complex substrate for TF interaction assays. Since the assembly system is derived from *Drosophila* preblastoderm embryos, the chromatin that arises has all hallmarks of pre-MBT embryos: it does not contain significant concentrations of endogenous TFs or components of the transcription machinery. The addition of defined TFs somewhat mimics aspects of zygotic genome activation (ZGA), when newly translated proteins initiate the complex genome expression program. The analysis of large compendia of potential TF binding sites in different genomic contexts allows statistical generalization, in addition to anecdotal illustration.

Our experimental system is well suited to discovering intrinsic protein-DNA interactions of well-characterised components. These may even include heterologous TFs of other species that bind short recognition motifs statistically distributed in *Drosophila* genomes. Characterising the intrinsic binding properties of TFs in a complex chromatin environment is important, even if TF binding specificities commonly observed *in vitro* do not explain physiological chromosome interactions (31).

Our focus has been on the DNA binding proteins that target the ‘male-specific-lethal’ (MSL) dosage compensation complex (DCC) exclusively to X chromosome-specific DNA elements to deploy the vital activation of transcription. *In vivo*, the MSL proteins do not bind autosomal DNA. We found that (1) MSL2, the DNA-binding subunit of the DCC, bears intrinsic sequence selectivity for a complex X chromosome-specific DNA signature, which we termed PionX sites (5); (2) MSL2 cooperates with the abundant GA-binding protein CLAMP to associate with *bona fide* HAS on the X chromosome *in vivo* (8). *In vitro* this cooperation results in the recruitment of MSL2 to hundreds of strong CLAMP binding sites on all chromosomes (8,11); (3) the GA-binding GAF competes with CLAMP/MSL2, preventing binding to a class of unphysiological sites (11). We now investigated the mechanism underlying the CLAMP-MSL2 cooperativity.

### Cooperativity turned into competition

We found that the cooperativity between CLAMP and MSL2 depends on the physical interaction between the two proteins. Although we detected a protein complex in solution, it is possible or even likely that the interaction is promoted by DNA (16). We expected that upon disruption of the TF interaction the direct cooperativity would turn into indirect, nucleosome-mediated cooperativity, the dominant mechanism of synergism between TFs (21,22,32). To our surprise, we found that under those conditions, the stronger GA-binder, CLAMP, competes with MSL2 for binding to composite sites. These sites are unusually long: MEME detects PWMs of 20 bp and more in peaks, so there should be space for two or more proteins to bind side by side. This is in line with the comprehensive study of Taipale and colleagues, who showed that often cooperating TFs bind composite sequence elements where individual TF recognition motifs overlap (16).

Since CLAMP binds degenerate GA repeats (27), it is possible that two CLAMP molecules can bind to these long sequences, or that single CLAMP molecules bind with variable translational positions, thus occluding GA sequences required for MSL2 interaction (see model in Figure 2E). Accordingly, MSL2 can resist competition at sites that contain the PionX signature, but will be competed away from sites of lower affinity that contain degenerate GA repeats (Figure 4B). In such a situation, the physical interaction between CLAMP ZF1 and the CBD of MSL2 may ensure that both proteins lock into a defined geometry that prevents competition.

### Relevance and limitation of intrinsic sequence specificity

MSL2 and CLAMP mutually cooperate to bind stably to X-chromosomal HAS *in vivo* (7,8). However, the chromosomal MSL2 binding profile only overlaps to a low extent with the pattern of MSL2-CLAMP binding *in vitro*. In some ways, this is not surprising given that MSL2 functions as a subunit of the large DCC, which contains other protein subunits as well as the long, noncoding *roX* RNA. Indeed, several studies concluded that the presence of *roX* RNA is required for faithful localization of MSL2 on the X chromosome (33,34). We found MSL2 delocalized to strong CLAMP binding sites on all chromosomes in the absence of *roX* RNA *in vivo* (6). Although the underlying mechanism is unknown, we note that *roX2* interacts with the C-terminus of MSL2, which also harbors the intrinsic GA-binding specificity of MSL2 as well as the CBD (35,36). We speculated that *roX* may modulate the DNA and protein interactions of MSL2, perhaps by restricting the cooperativity with CLAMP (6). Remarkably, Larschan and colleagues recently observed that at nuclear division cycle 13, prior to the expression of *roX2* and the assembly of a mature DCC, MSL2 and CLAMP localize to CLAMP binding sites on all chromosomes (37). The binding profile we describe in reconstituted preblastoderm chromatin may reflect some of these earliest MSL2 interactions with chromosomes prior to *roX2* expression.

### Identification of an expanded set of X-chromosomal MSL2 binding sites

Taking advantage of the superior resolution and signal/noise output of C&R analyses we determined MSL2 binding sites in S2 cells. We obtained more that 700 distinct peaks of MSL2 interaction, all of which were localized on the X chromosome. Stratifying the peaks according to their signal strength reveals a hierarchy of sequences. Motifs with 5’ extension, resembling the PionX motif ranged among the sites with highest affinity, followed by sites of intermediate affinity that display the MRE consensus sequence. Weaker binding sites contain degenerate MRE motifs dominated by GA repeats. These data support earlier conclusions from low-resolution studies on polytene chromosomes about the existence of a hierarchy of binding sites, where sites of higher affinity are primary attractants for MSL proteins and sites of lower affinity profit from local TF enrichment (24,30).

### Is there a defined anatomy of high affinity sites for the MSL2?

The X-chromosomal HAS or ‘chromosome entry sites’ were initially defined at low resolution on polytene chromosomes. With the advent of the ChIP-seq methodology it became clear that the peaks of DCC binding are several hundred base pairs long and often contain two or more MRE sequences (38,39). MEME analysis often focuses only on the strongest MRE within a larger ChIP-seq peak. Intuitively, loci with multiple MREs may attract more MSL2 than sites with a single MRE. It is also conceivable that higher affinity sites have a defined arrangement of sites, which may facilitate cooperative interactions between MSL complexes bound to different elements. However, to our knowledge these questions have not been systematically investigated.

With respect to the diversity of MRE number and arrangement, which may or may not include PionX motifs, the intrinsic binding sites obtained *in vitro* and the physiological X chromosomal binding sites appear very similar. In both cases, we found no strong correlation between the number, length or arrangement of MRE sequences with ChIP-seq peak intensity. We conclude that the binding strength is mainly determined by the affinity of MSL2 for individual MRE motifs, where the cooperativity with CLAMP plays an important role. The ‘dominant’ binding event can be inferred from our EChO analyses. MSL2 appears particularly dependent on this cooperation, since embryonic nuclei only contain very limited amounts of MSL2, barely enough to distribute one copy for each binding site mapped by C&R (40). Quinn *et al*. suggested that MREs originally involved from pyrimidine-rich splicing enhancers in introns and were not under selection for precise location, but rather for proximity to genes (41). Our finding that MSL2/CLAMP binding does not require a defined architecture of binding sites resonates with the observation that defined spacing and orientation of multiple binding sites are commonly not needed to assure integration of different TF input (17).

In conclusion, our analyses show that the direct contact between MSL2 and CLAMP ensures cooperativity and avoids competition between the two proteins at individual MREs. Remarkably, *in vivo* this intrinsic mechanism only operates at X chromosomal sites where the DCC is functional. Future research will focus on finding the molecular mechanism that restricts this cooperative binding to the X chromosome.

## Methods

### Co-Immunoprecipitation

The protocol was adapted from (8). Briefly, we mixed 50 nM of recombinant MSL2-FLAG and CLAMP-FLAG in 100 µl DREX, and incubated the mixture with end-over-end rotation for 30 minutes at room temperature (RT). Next, we added the mixture to 20 µl of antibody-coupled protein-AG beads, which had been pre-coupled with MSL2 antibody for 3 hours at 4°C with end-over-end rotation. The beads were then washed three times with 1 ml of EX50 buffer (10 mM Hepes/KOH pH 7.6, 50 mM KCI, 1.5 mM MgCl2, 50 µM ZnCl_2_, 10% glycerol, 1 mM DTT, and 1x cOmplete EDTA-free Proteinase inhibitor cocktail (PIC, Roche)), followed by incubation with the IP reaction for 30 minutes at room temperature. After incubation, the beads were washed three more times with 1 ml of EX50 buffer, resuspended in Laemmli buffer, and processed for western blotting.

### Western Blots

Samples were denatured with Laemmli buffer at 95°C for 5 min. Then samples were electrophoresed on 4-20% SDS ServaGel TGPrime (Serva) for 1 h at 100 V. Proteins were transferred to AmershamTM ProtranTM 0.45 µM Nitrocellulose Blotting Membrane for 1 h at 400 mA. All following incubations were for 1 h at RT. Membranes were blocked with 5% BSA. The membrane was incubated with primary antibody in TBS-T, washed thrice with TBS-T and incubated with secondary antibody. Images were taken using the LICOR Odyssey CLx.

### DNA purification

Genomic DNA (gDNA) was obtained from male BG3-c2 cells (Drosophila Genomics Resource Center) (42). They were cultured at 26°C in Schneider’s Drosophila Medium (GIBCO) with 10% fetal calf serum (FCS), Penicillin-Streptomycin and 10 mg/ml human insulin. The DNA of 10^7^ cells was purified using the Blood & Cell Culture DNA Midi Kit (Qiagen) following the supplier’s protocol. The DNA was dissolved in EDTA-free 10 mM Tris-NaCl pH 8. Concentrations were determined using Qubit (Thermo Fisher).

### Preparation of prebastoderm embryo chromatin assembly extract (DREX)

Preblastoderm embryos were collected within 90 min after egg laying, as previously described (12). Briefly, settled embryos (50 ml) were dechorionated in 200 ml of embryo wash buffer (EW) (0.7% NaCl and 0.04% Triton X-100), followed by treatment with 13% sodium hypochlorite (VWR) for 3 min at RT with constant stirring. After rinsing the embryos for 5 min with cold water, they were transferred to a glass cylinder containing EW and allowed to settle. The settled embryos were then washed successively with 0.7% NaCl and extract buffer, consisting of 10 mM Hepes/KOH pH 7.6, 10 mM KCI, 1.5 mM MgCl2, 10% glycerol, 1 mM DTT, and 1x cOmplete EDTA-free Proteinase inhibitor cocktail (PIC, Roche).

The embryos were homogenized using a homogenplus homogenizer (Schuett-Biotec) with one stroke at 3000 rpm and 10 strokes at 1500 rpm after which the MgCl_2_ concentration of the homogenate was adjusted to 5 mM (final concentration) and centrifuged at 27,000 g for 15 min at 4°C. The lipid layer was discarded, and the supernatant was further centrifuged at 245,000 g for 2 h at 4°C. The clear extract was collected using a syringe, while the lipid layer and pellet were left behind. To avoid chelation of Zn, EDTA was excluded from all steps of the purification. Extracts were stored in 200 µl aliquots at −80°C after shock freezing in liquid nitrogen. Extracts were only thawed once before use.

### Chromatin assembly

One µg of genomic DNA was assembled into chromatin by adding 15 µl 10x McNAP buffer (0.3 M creatine phosphate, 30 mM ATP, 3 mM MgCl_2_ 1 mM DTT, 10 ng/µl creatine phosphokinase), 100 µl DREX extract and EX50 buffer (10 mM Hepes/KOH pH 7.6, 50 mM KCI, 1.5 mM MgCl_2_, 50 µM ZnCl_2_, 10% glycerol, 1 mM DTT, 1x PIC) up to a final volume of 150 µl. The amounts of extract necessary were determined empirically for each batch. Assembly took place at 26°C for 4 h.

### MNase digestions

One µg of DNA assembled into chromatin in 150 µl was digested with MNase by adding 200 µl MNase digestion solution (186 µl EX50, 10 µl 1 M CaCl_2_ and 4 µl MNase solution 333 U/µl). At times 15, 30 and 120 sec, 110 µl were transferred to tubes containing to 40 µl 100 mM EDTA solution each to stop the digest. Two µl glycogen (10 mg/ml) and 150 µl 7.5 M ammonium acetate were added and samples were mixed. Then, 880 µl of 100% ethanol was added and samples were vortexed vigorously and cooled at −20°C for 10 min. After centrifugation at 21,000 g for 15 min at 4°C, the supernatant was removed and pellets were washed with icecold 70% ethanol. After pelleting the DNA again at 21,000 g for 5 min at 4°C, it was dissolved in 8 µl 10 mM TE buffer and 2 µl Orange-G loading dye. Samples were separated on a 2% agarose gel, stained with ethidium bromide and imaged using the Quantum ST-4 from PeqLab to control for proper chromatin assembly. Samples used for later ChIP-seq experiments were always digested for 2 min.

### Baculovirus infections

Sf21 cell cultures at 10^6^ cells/ml (2.5 ×10^8^ cells) were infected 1:1000 (v/v) with baculovirus, expressing the respective FLAG tagged proteins as described in (43). After 72 h, cells were harvested and washed once in PBS, frozen in liquid nitrogen and stored at −80°C.

### Protein purification

EDTA was excluded from all steps of the purification to avoid chelation of Zn in later experiments.

MSL2-FLAG: Sf21 cell pellets were rapidly thawed and resuspended in ice-cold Lysis buffer (300 mM KCl, 50 mM Hepes/KOH pH 7.6, 5% glycerol, 0.05% NP-40, 1 mM MgCl_2_, 50 µM ZnCl_2_, 1 mM DTT, 1x PIC). 25 ml buffer was added to the cell pellet (2.5 ×10^8^ cells). After 15 min incubation on ice, the suspension was sonicated (5 × 10 sec pulses, 20 sec break, 20% amplitude, Branson sonifier model 250-D) and centrifuged for 45 min at 30,000 g at 4°C. The soluble protein fraction was incubated with Lysis buffe-equilibrated FLAG beads (Anti-FLAG M2 Agarose, Sigma) for 3 h at 4°C on a rotating wheel. 0.5 ml beads were used per 2.5 ×10^8^ cells. The beads were washed twice with 10 ml ice-cold Lysis buffer, twice with 10 ml Wash buffer (Lysis buffer, but with 1 M KCl and 1% NP-40) and twice with 10 ml Elution buffer (Lysis buffer, but with 100 mM KCl). The FLAG-tagged MSL proteins were eluted for 3 h at 4°C on a rotating wheel in the presence of 0.5 mg/ml FLAG-Peptide (Sigma) in 1 ml Elution buffer. Purified proteins were then rapidly frozen in liquid nitrogen and finally stored at −80°C. Protein concentrations were determined via SDS–PAGE and Coomassie staining using BSA (NewEngland Biolabs) as a standard. Cloning for the MSL2 expression construct is described by (43).

CLAMP-FLAG and CLAMP-ΔZF1-FLAG: Sf21 cell pellets were rapidly thawed and resuspended in 1 ml Buffer C per 10 mL of culture (50 mM HEPES pH7.6, 1 M KCl, 1 mM MgCl_2_, 5% (v/v) glycerol, 0.05% NP-40, 50 μM ZnCl_2_, 375 mM L-Arginine (according to (44)) supplemented with 0.5 mM TCEP and 1x PIC. After 15 min incubation on ice, the suspension was sonicated (5×10 sec pulses, 20 sec break, 20% amplitude, Branson sonifier model 250-D). The extract was adjusted with Buffer C containing 1x PIC to 2 ml per 10 mL of culture and supplemented with 0.1% (v/v) polyethyleneimine by adding 2% (v/v) polyethyleneimine (neutralized with HCl to pH 7.0) drop-by-drop while string in an ice bath [according to (45)] and then centrifuged for 45 min at 30,000 g at 4°C. The soluble protein fraction was incubated with Buffer C-equilibrated FLAG beads (Anti-FLAG M2 Agarose, Sigma) for 3 h at 4°C on a rotating wheel. 0.5 ml beads were used per 2.5 ×10^8^ cells. Beads were pelleted at 4°C for 5 min at 500 g and supernatant was removed. Beads were washed 5 times with 20 bed volumes of Buffer C. The FLAG-tagged CLAMP proteins were eluted for 3 h at 4°C on a rotating wheel in the presence of 0.5 mg/ml FLAG-Peptide (Sigma) in 1 ml Buffer C containing 1x PIC. Purified proteins were then rapidly frozen in liquid nitrogen and finally stored at −80°C. Protein concentrations were determined via SDS–PAGE and Coomassie staining using BSA (New England Biolabs) as a standard. Cloning for the CLAMP constructs is described by (8).

### Antibodies

The following antibodies were used: rabbit α-MSL2 (43); guinea-pig α-MSL2 (8); rabbit αCLAMP (8); IRDye 680RD goat α-rabbit (Licor 925-68071); IRDye 800CW goat α-guinea-pig (Licor 926-32211).

### ChIP-seq

Recombinant Proteins were added to 1 µg of assembled chromatin as described in ‘chromatin assembly’ and were allowed to bind for 1 h. Samples were crosslinked by adding formaldehyde (Thermo Fisher Scientific, Ref 28908) up to 0.1% final concentration for 10 min and then quenched by 125 mM glycine for 5 min. Samples were digested by MNase as described above for 2 min. After adding 1x RIPA buffer up to 500 µl, samples were precleared on a rotating wheel with 20 µl buffer-equilibrated protein-AG beads per 1 µg chromatin for 1 h at 4°C. Supernatant was collected and 2 µl of purified antibody was added to the precleared sample and let to bind overnight at 4°C on a rotating wheel. Then samples were bound to freshly washed protein AG beads for 3 h. Samples were washed 4 times for 5 min with 1 ml of 1x RIPA buffer per sample (1 µg chromatin on 20 µl beads). Then the beads were suspended in 100 µl 1x TE buffer and de-crosslinked overnight at 65°C while shaking. Samples were then digested with 10 µg RNAse A for 30 min at 37°C and 100 µg proteinase K at 56°C for 1h. beads were pelleted at 1000 g for 1 min and supernatant was transferred to a fresh tube for purification. DNA was purified by two extractions with Phenol:Chloroform:Isoamyl-alcohol (25:24:1, Sigma Aldrich) and precipitated by adding it to 2 µl of glycogen, 0.1 × volume 3 M sodium acetate. Then 2.5 × volume 100% ethanol was added and samples cooled at −20°C for 15 min and pelleted in a tabletop centrifuge. The DNA was washed once with 70% ethanol and dissolved in EDTA-free 10 mM Tris/NaCl, pH 8. Concentrations were determined using Qubit (Thermo Fisher).

For correlation analyses between replicates, see Figure S2.

### CUT&RUN

CUT&RUN was performed as described in (29). Briefly, 10^6^ S2 cells were harvested, washed and bound to pre-activated Concanavalin A-coated magnetic beads, in wash buffer (20 mM HEPES, pH7.5, 150 mM NaCl, 0.5 mM spermidine and 1X cOmplete EDTA-free PIC). The cells were then permeabilized in Antibody buffer containing 0.05% digitonin (Dig-Wash) and 2 mM EDTA and incubated with rabbit α-MSL2 serum or pre-immune serum, diluted 1/500, at 4°C overnight on a nutator. After antibody incubation, the beads were washed thrice with DigWash buffer and incubated with pAG/MNase at a concentration of 700 ng/ml, for 1 h at 4°C. After 2 washes in Dig-Wash buffer the beads were resuspended in 100 μl of Dig-Wash buffer and chromatin was digested, by addition of 100 mM CaCl_2_, for 30 min at 0°C. The digestion reaction was stopped by addition of 100 μl 2X STOP buffer (340 mM NaCl, 20 mM EDTA, 4 mM EGTA, 0,05% Digitonin, 50 µg/ml glycogen, 100 µg/ml RNase A). The digested chromatin fragments were released by incubating the beads for 30 min at 37°C. The beads were then placed on a magnet stand and the liquid containing the digested chromatin was transferred to a fresh tube. Total DNA was extracted using the Phenol/Chloroform protocol. Concentrations were determined using Qubit (Thermo Fisher).

For correlation analyses between replicates, see Figure S2.

### Library preparation and sequencing

For ChIP-seq samples, libraries were prepared using NEBNext Ultra II DNA Library (New England Biolabs) according to manufacturer’s instructions. For CUT&RUN, sequencing libraries were prepared as described in (46,47). All libraries were sequenced on an Illumina NextSeq1000 sequencer. About 20 million paired-end reads were sequenced per sample for each of the ChIP replicates. Replicates were performed using a separate batch of purified proteins and DREX extracts. For CUT&RUN about 5 million paired-end reads were sequenced per sample for each of the biological replicates. Base calling was performed by Illumina’s RTA software, version 1.18.66.3.

## Data analysis

### Read processing

Sequence reads were Demultiplexed by JE demultiplexer (48) using the barcodes from the Illumina Index read files. Demultiplexed files were aligned to the *Drosophila melanogaster* release 6 reference genome (BDGP6) using Bowtie2 (49) version 2.2.9. (parameter “--end-toend −-very-sensitive −-no-unal −-no-mixed −-no-discordant −X 400”) and filtered for quality using samtools 1.6 (50) with a MAPQ score cutoff of −q 2.

### Replicate Correlation

Replicate correlation was determined by first searching the dm6 genome for 5000 best hits of the GAGAGA MRE-motif by FIMO (MSL2-ChIP n=3, CLAMP-ChIP n=2). Then each replicate was down-sampled to receive the same number of reads per replicate, and reads per motif were counted and plotted against each other. If replicates were sufficiently similar, the sampled reads were merged and used for further analysis. This avoids normalization against an input and retains individual read information. For correlation analyses between replicates, see Figure S2.

### Peak calling

Peaks were called using Homer (51) version 4.9.1 calling the functions makeTagDirectory (parameters −single −fragLength 150) and findPeaks (parameters −style factor −size 150 −F 6) using the corresponding negative samples in which the IP was done without adding the respective protein as control. We called peaks against a negative control (IP in the absence of added protein) where possible, as this allows us to account for antibody cross reactivity in immunoprecipitations. Peak calling was done with the summarized replicates for each sample and the control, resulting in more robust peaks through the additional coverage used. HAS and PionX regions and motifs were used as defined by (5) with 309 HAS and 56 PionX in total.

### De novo motif discovery

Enriched motifs in peak region were discovered using MEME (52) (version 4.11.4 (parameters −mod anr −dna −revcomp −nmotifs 2). Before analysis the peaks were resized to 200 bp to include 25 bp of sequences directly bordering the peaks.

### Find Motif occurrences

Motif search using position weight matrixes from MEME in peak regions on the genome was performed with FIMO (53) version 5.0.2.

### Browser profiles

Browser profiles were generated using UCSCutils (http://genome.ucsc.edu.) version 3.4.1. calling the function makeUCSCfile using the summarized sample replicates Tag Directories, also used for the peak calling, and were normalized against the control. Values are fold change over control. Profiles were visualized using the IGV software (54).

### Data Analysis and plotting

Data Analysis was conducted in R (55), using the tidyverse libraries (56).

### Heatmaps and cumulative plots

Heatmaps were made using the R library “Complexheatmaps” (57) by cutting windows of 2000 bp around sites of interest of the calculated coverages normalized against a control if applicable and aligning them. The cumulative plots are made by calculating the mean of each column. Window identities are retained in the data and used for the annotation by overlapping them with the known HAS or the X chromosome.

### Fragment size analysis

Plots depicting individual fragments at each peak and annotation of motifs within peaks was done by custom code using the libraries zoo (58), rtracklayer (59) and ggpmisc.

### Alphafold

Protein structures were predicted using ColabFold (60) which uses the Alphafold2 algorithm (61).

### Data availability

For the in vitro reconstitution assays the raw sequencing files in fastq format and the summarized genome browser tracks in bigwig format are available in the GEO database at GSE224981.

To review GEO accession GSE224981:

Go to https://www.ncbi.nlm.nih.gov/geo/query/acc.cgi?acc=GSE224981

Enter token wpsvyscupdutdcv into the box.

For the Cut and Run experiments the raw sequencing files in fastq format and the summarized genome browser tracks in bigwig format are available in the GEO database at GSE228935.

To review GEO accession GSE228935:

Go to https://www.ncbi.nlm.nih.gov/geo/query/acc.cgi?&acc=GSE228935 Enter token khejieswvzkrpuf into the box.

Custom code can be found as Zenodo repository und number 7816989 DOI: 10.5281/zenodo.7816989

## Acknowledgments

We thank S. Krebs from the LAFUGA Genomics Facility for next-generation sequencing. We thank T. Straub and T. Schauer (BMC Bioinformatic Core Facility) for scripts and advice and all members of the Becker lab for discussions throughout the project. We thank E. Larschan for the generous gift of α-CLAMP antibody. This work was supported by grant Be1140/8-1 of the German Research Council (DFG) to PBB.

## Author Contributions

NE and PBB conceived experiments; NE and FG performed experiments; SK cloned baculovirus expression constructs and carried out viral infections; ACS prepared the sequencing libraries and optimized the CUT&RUN protocol; NE analyzed data; PBB provided feedback and supervision; NE and PBB wrote the manuscript; PBB secured funding.

