## Supplement_figure_1 for "Physical interaction between MSL2 and CLAMP assures direct co-operativity and prevents competition at composite binding sites"

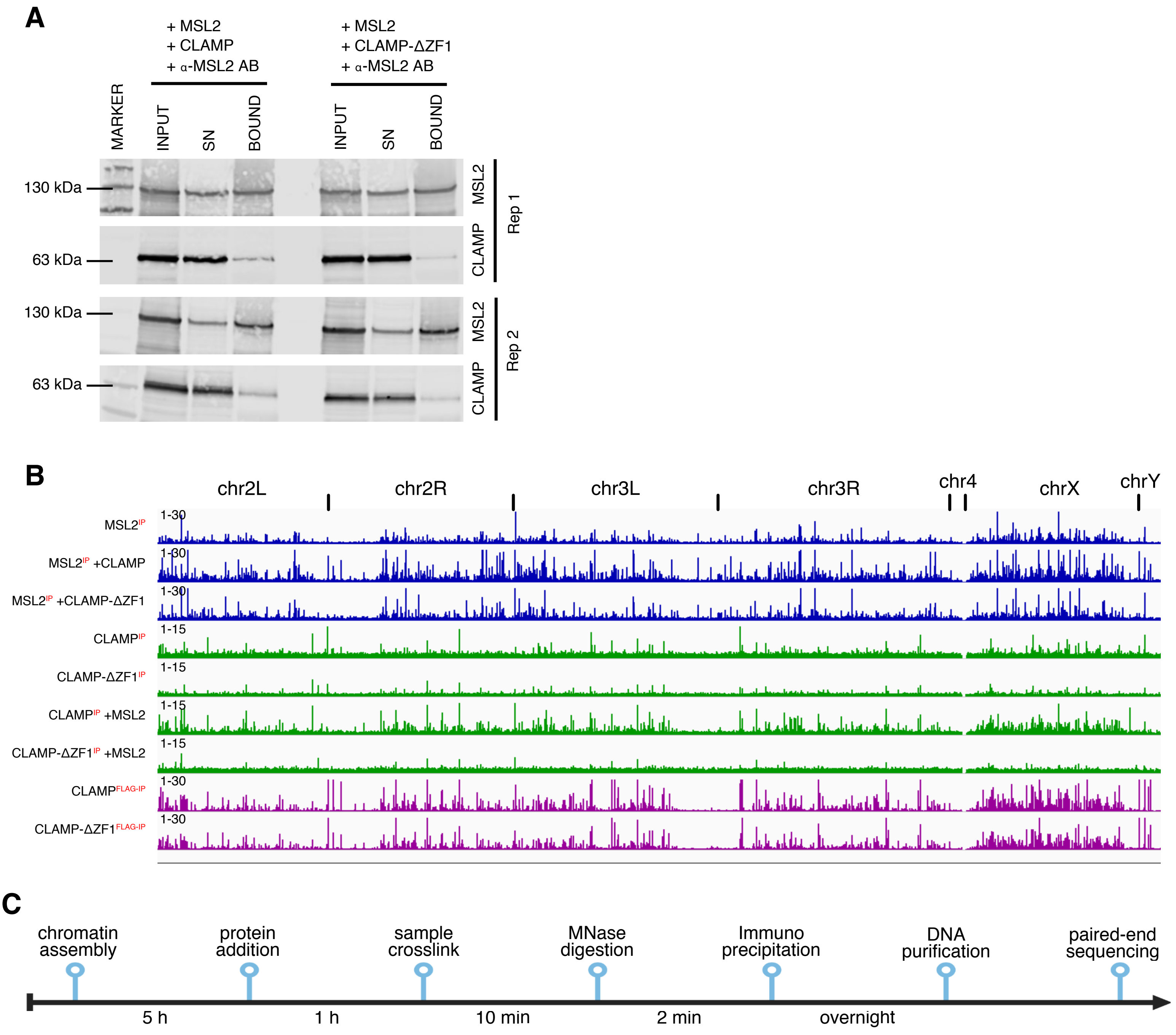


Figure S1

(A) Two replicate Western blots of co-immunoprecipitation of MSL2 and CLAMP. For legend see Figure 1, where one of the plots is shown.

(B) Genome browser profile all summarized biological replicates showing binding of the protein marked with 'IP' along the entire genome, determined by ChIP-seq (MSL2-IP n=3, CLAMP IP n=2). Maximum values for each window are shown and potential binding sites (MREs) and chromosomes are annotated.

(C) Time line of the chromatin reconstitution and in vitro ChIP experiments.
