## Supplement figure_2 for "Physical interaction between MSL2 and CLAMP assures direct co-operativity and prevents competition at composite binding sites"

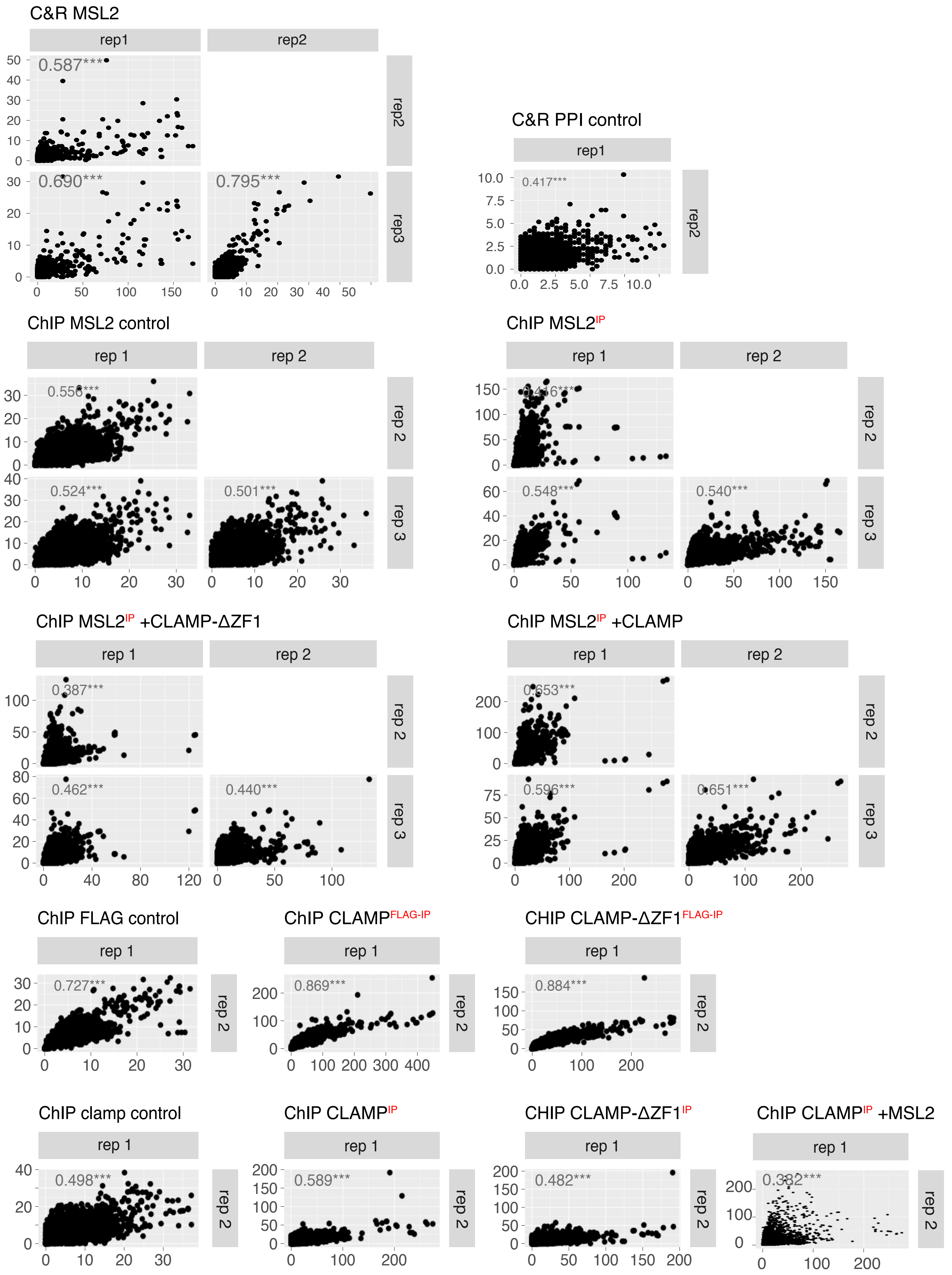


Figure S2: (A) Replication Correlation analysis for all sequencing samples used in the study. For in vitro samples, chromatin was assembled in the presence and absence of the annotated proteins and DNA binding of the protein marked with 'IP' was determined by ChIP-seq. For in vivo samples, binding was determined by CUT&RUN. Reads were sampled to get the same read count for each replicate and quantified at 5000 computationally determined potential binding sites. Sample names and R value are annotated and p values were coded as asterixis in the following manner: p= 0.05*, 0.01**, 0.001***. Correlating sampled reads were then summed up for further analysis.
